# Circuit-level Mechanisms of EtOH-dependent dopamine release

**DOI:** 10.1101/119859

**Authors:** Matteo di Volo, Ekaterina O. Morozova, Christopher C. Lapish, Alexey Kuznetsov, Boris Gutkin

**Author notes:** Corresponding author: Matteo di Volo. The authors declare no competing financial interests.

## Abstract

Alcoholism is the third leading cause of preventable mortality in the world. In the last decades a large body of experimental data has paved the way to a clearer knowledge of the specific molecular targets through which ethanol (EtOH) acts on brain circuits. Yet how these multiple mechanisms play together to result in a dysregulated dopamine (DA) release under alcohol influence remains unclear. In this manuscript, we delineate potential circuit-level mechanisms responsible for EtOH-dependent increase and dysregulation of DA release from the ventral tegmental area (VTA) into nucleus accumbens (Nac). For this purpose, we build a circuit model of the VTA composed of DA and GABAergic neurons, that integrate external Glutamatergic (Glu) inputs to result in DA release. In particular, we reproduced a non-monotonic dose dependence of DA neurons firing activity on EtOH: an increase in firing at small to intermediate doses and a drop below baseline (alcohol-free) levels at high EtOH concentrations. Our simulations predict that a certain level of synchrony is necessary for the firing rate increase produced by EtOH. Moreover, EtOH’s effect on the DA neuron’s firing rate and, consequently, DA release can reverse depending on the average activity level of the Glu afferents to VTA. Further, we propose a mechanism for emergence of transient (phasic) DA peaks and the increase in their frequency in EtOH. Phasic DA transients result from DA neuron population bursts, and these bursts are enhanced in EtOH. These results suggest the role of synchrony and average activity level of Glu afferents to VTA in shaping the phasic and tonic DA release under the acute influence of EtOH and in normal conditions.

## Introduction

Alcoholism is the third leading cause of preventable mortality in the world (Mokdad et al. 2004) and causes 2.5 million deaths per year. In the last decades a large body of experimental data has paved the way to a clearer knowledge of the specific molecular targets through which ethanol (EtOH) acts on brain circuits (Morikawa et al. 2010).

Multiple observations suggest that the behaviors associated with EtOH (seeking, consumption, etc) are at least partially dependent on the mesolimbic Dopamine (DA) system, e.g. it has been shown that DA_1_ receptor blockade in NAc suppresess EtOH consumption (Dyr et al. 1993, and Samson et al. 1993 and Gonzales et al. 2004). Acute EtOH has been largely shown to produce rapid increase of DA concentration in NAc, a brain region crucial for cognitive processing of reward and reinforcement learning, and hence with a significant role in addiction (Yim, H. J et al. 2000). Such an effect is thought to be due to the alcohol influence on the DA neurons in the Ventral Tegmental Area (VTA) that project directly to the NAc (Pierce et al. 2006). More specifically, after acute alcohol injection, NAc DA shows a significantly more frequent (as compared to alcohol-free control) high-amplitude transient peaks of concentration (Covey et al. 2014). Such changes in the dopamine release pattern stems from the effect of alcohol on the firing activity of DA neurons in the VTA. Alcohol, by modifying their dynamics and computational properties, produces a dysregulation in DA levels in NAc characterized by high-amplitude fluctuations. In turn these phasic fluctuations may lead to increased motivational valence for drug-seeking.

In this manuscript we delineate potential circuit-level mechanisms responsible for EtOH-dependent increase and dysregulation of DA release to NAc. For this purpose we build a circuit model of the VTA composed of DA and GABAergic neurons, that integrate external Glutamatergic inputs to result in dopamine released by DAergic neurons at their target sites.

In particular, we concentrate on the way EtOH shows a non-monotonic effect on DA neurons firing activity: an increase in the firing rate at relatively small doses and a drop below baseline (alcohol-free) levels at high EtOH concentrations. Furthermore, we propose a mechanism for the increase in DA neurons phasic activity due to EtOH. Finally, we focus on the effect of EtOH on DA concentration dynamics, by analyzing the emergence of high-amplitude transient peaks produced by activity of a heterogeneous population of DAergic neurons in the VTA.

We investigate how these features can be reproduced by considering only EtOH effects on DAergic neuron proprieties (input conductances and intrinsic currents), not considering the effects of EtOH on activity patterns of neural populations projecting to DAergic neurons in the VTA (eg. PFC). The latter is a subject of a separate study.

Following experimental observations, we model major targets EtOH on DAergic neurons (Morikawa et al. 2010), i.e. on their intrinsic properties and on the synaptic receptors activated by the external inputs. Regarding the first one, we calibrate the model according to *in vitro* experiments. We thus calibrate EtOH effects on the intrinsic conductances (HCN and GIRK) according to measurements on EtOH action on DAergic neuron firing rate. Regarding DA neuron ligand-gated ion channels responsible for integrating external inputs, we simulate ETOH effect by a calibrated increase in the AMPA-to-NMDA ratio and in the GABA conductance as suggested in Morikawa et al. (2010).

All together, this extrapolation from experimental evidences will permit us to exploit a circuit model computing EtOH effects on VTA at various concentration. In particular, we will describe the way VTA processes, under different EtOH doses, external glutamatergic (Glu) inputs for dopamine release.

In the next section, we briefly describe the circuitry we model for DAergic neuron activity simulation and EtOH effects. The Results section is then divided in three parts. In part 1 we report simulations showing the model’s capability to reproduce a inverted U-shape of EtOH effect on DA neuron activity observed in *in vivo* measurements: an increase in VTA DAergic neuron average firing rate at relatively low ETOH concnetrations, followed by a decrease at high concnetrations. Interestingly, our simluations predict that the EtOH effects on DA neuron bursting require Glu inputs to DA neurons to be synchronized so as to produce a temporally varying levels of DA neuron activity. Our modelling results show that Glu inputs to the VTA need to be synchronized and variable in time in order to produce increased DA cell bursting due to EtOH intake. Synchronization in AMPA inputs generates higher values of AMPA EPSC. Nevertheless, the effect here described does not depend on the amount of AMPA current flowing in DA neurons but on the variability of such current in time as due to synchronous inputs. In order to emphasize this effect we analyse a reduced model where we increase input synchronization but we change AMPA conductance in order to keep fixed the average amount of AMPA current flowing in DA neuron. We show that the olnly increase in synchronization and variability in AMPA inputs yields a transition to high rate phasic activity in DA neurons.

We describe how a greater hyperpolarizing GABA tone (as driven by EtOH) can increase DA neuron bursting (as measured by percentage of spikes within bursts, SWB). In Section 2 we report how EtOH effects depend on the levels of Glu input activity. We show that, while low and moderate frequency Glu input causes EtOH to have an excitatory effect on the DA neuron, during high firing rate Glu input, EtOH becomes inhibitory. In fact, in this setup DAergic neuron is hyperexcited by EtOH and its firing is blocked. Furthermore, we show that the presence of EtOH changes the temporal dependence of the VTA response to its inputs: over a critical EtOH concentration, DAergic neuron activity transits from being *in phase* to *antiphase* with respect to Glu inputs. In part 3, we consider a heterogeneous population of DAergic neurons, that permits us to generate DA release from VTA. We report that EtOH has an inverted U-shaped effect on the average DA concentration: an increase followed by a decrease at high EtOH doses. We then show that EtOH is able to increase transient fluctuations in DA concentration and describe a mechanism for this increase via synchronization of the VTA DAergic neurons.

## Results

### EtOH & VTA input-output processing: overview of the circuit model

The computational model takes into account the circuitry involved in DAergic neurons activity in the VTA and its modifications due to EtOH. We model VTA as composed of a network of electrically (gap-junction) connected and hererogeneous GABA neurons that project to a heterogeneous population of DAergic neurons (Margolis et al. 2012 and Roeper 2013). This feed-forward inhibitory circuit is suggested by experimental evidences (Steffeensen et al. 1998). Furthermore, the VTA receives excitatory inputs from cortical regions, specifically excitatory cells in the PFC (Carr et al. 2000). Accordingly, for the inputs, we use a putative model generating Poisson-distributed glutamatergic spike trains with a degree of synchronization between them that we define as a free parameter. In order to construct such synchrony, we force a certain fraction f_s_ of such neurons to fire almost simultaneously (their firing events are randomly distributed in a 5ms time window) in certain time windows (see method for more details and Fig.1A as an example). These inputs, projecting to the VTA, are thus characterized by two main parameters: the average firing rate (common to all units) ν_Glu_ and their level of variability taking into account synchronization. This excitatory activity furnishes inputs to the VTA through AMPA and NMDA receptors located on both GABA and DA neurons. The latter receive also an inhibitory input through GABA_A_ receptors (see method section). In Fig.1A we show a cartoon of such circuitry.

**Figure 1:**
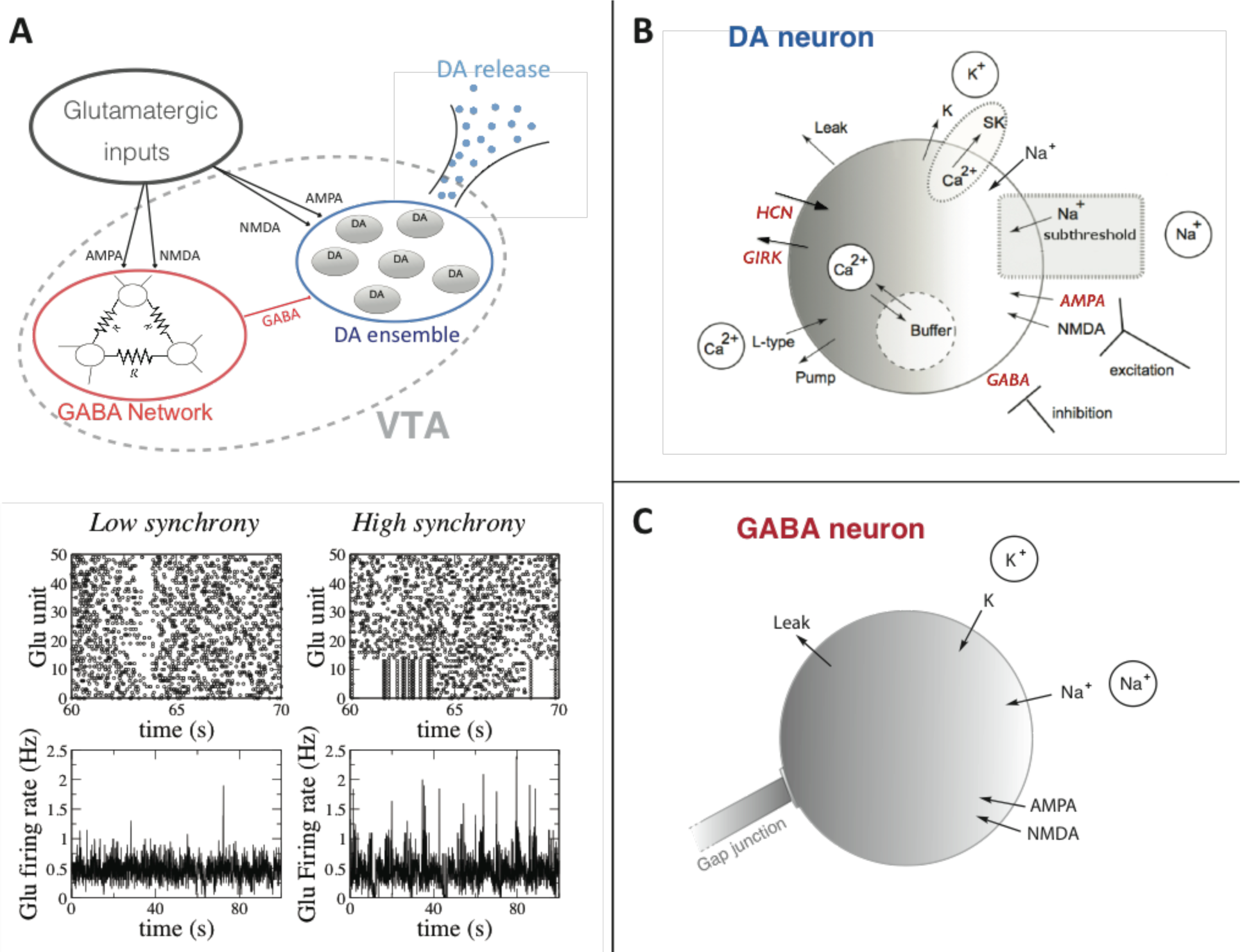
A cartoon of the Glu-VTA-DA circuit model. A) On the top, a population of Glutamatergic neurons send inputs processed by the VTA, which is composed of a population of electrically connected GABA neurons and a heterogeneous population of DAergic neurons. On the bottom, an example of Glu units firing pattern in the asynchronous and quasi-synchronous (f_s_ 16%) case. The Glu firing rate has been calculated through a normalized spike count of the population in a time bin of 10ms. B) Schematic view of an individual DAergic neuron model. In red we show the channels and conductances affected by EtOH. DA release from the DAergic neurons is computed through a synaptic release model. C)) Schematic view of an individual GABAergic neuron model.

The cascade of excitatory inputs is processed by the VTA local circuit, whose output (DA release) is computed through a synaptic DA release model developed by Wightemann et al. (1990), thus completing the structure of Glu-VTA-DA ouflow circuit model.

On the bottom of Fig1A we show an example of Glu units firing pattern (asynchronous and quasi-synchronous) with the relative measure of the population firing rate (normalized spike count) showing higher fluctuations in the quasi-synchronous case. In panels B and C we show the model of DA and GABA neurons, described in detail in the method section.

We will investigate how EtOH, through its action at the level of VTA, affects the input-output processing in this nucleus. For this purpose, we model the two most well-known classes of EtOH effects at the level of DAergic neurons. We include (1) the synaptic effects of EtOH, the increase in the GABA conductance as well as in the ratio of AMPA and NMDA synaptic currents in DA neurons, and (2) the DA neuron intrinsic excitability change due to increased HCN and GIRK channel conductances (see method section).

## 1. Impact of input synchronization on EtOH-evoked modification of DA neuron activity

In this section we describe the effects of various concentration of EtOH on the firing activity of a single DAergic neuron. In particular, we describe how EtOH effects are modified in two different Glu input states. In the first case we consider asynchronous inputs. These input patterns (Glu firing rate as in Fig.1) are characterized by a relatively low variability (f_s_=0%, the coefficient of variation of the Glu firing rate, as reported in Fig 1A is around 0.4). In the second case, we constrain a certain fraction of inputs to be synchronous (equal to f_s_=14%, the coefficient of variation of Glu firing rate, as reported in Fig 1A is around 0.65), obtaining input with a high degree of temporal variation.

### DAergic neuron firing rate shows U-shaped dependence on EtOH concentration

We first investigated effects of EtOH in our model on the firing rate of the target DA neuron. Our simulations show that under both the asynchronous (Fig2 left column) and partially synchronized (Fig2, right column) glutamatergic input, EtOH has a dual effect on the DA cell firing rate. In fact, for both cases, relatively low doses of EtOH progressively increase the firing rate up to a maximum, with a reversal of this effect at higher doses, and even a depression with respect to control at maximal doses. Thus the EtOH effect on the firing rate shows an inverted U-shaped curve. By carefully analyzing our model, we can explain this U-shaped response by two complementary mechanisms depending on endogenous cell excitability and glutamamtergic synaptic transmission in the first case, and dose-delayed GABA response in the second. First, at sufficiently low EtOH concentrations, the increase in AMPA and HCN channel conductance by EtOH produces an increase in DA neuron average firing rate (0-2 g/Kg). At these doses GABA conductance is not highly affected by EtOH. On the contrary, at high EtOH concentrations (e.g. 3g/Kg), GABA conductance on the DA neuron becomes sufficiently high to decrease the DA firing rate and bring it to smaller values compared to the baseline (no EtOH). We thus surmise, that for the effects on the firing rate, the EtOH effects on single DA neuron properties are sufficient to account for the inverted u-shaped effects observed in in-vivo experimental measurements (Mereu et al. 1984). This effect is observed regardless of the synchronization level of Glutamatergic inputs.

**Figure 2:**
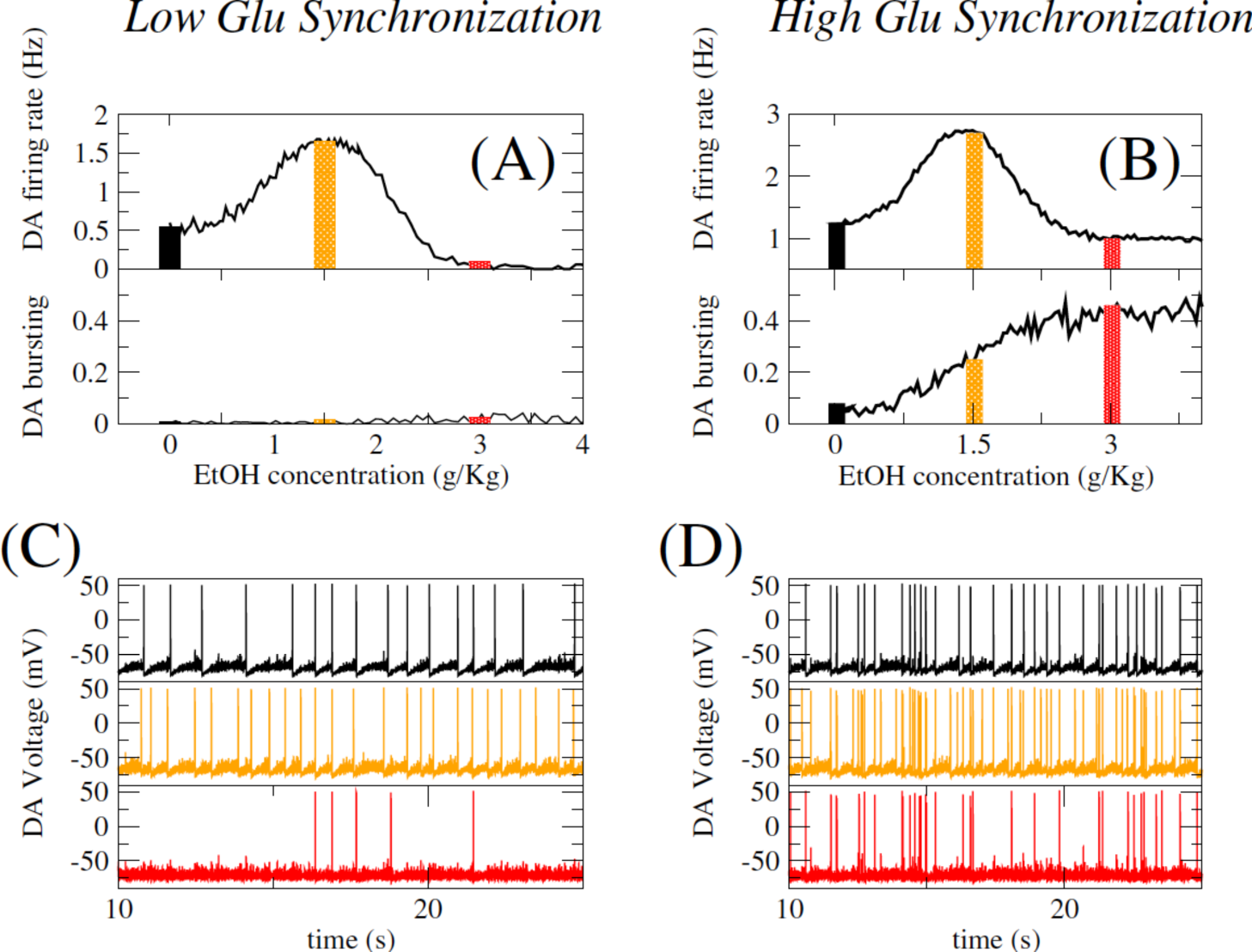
If combined with Glu input variability, EtOH boosts DA neuron activity and burstiness. On the left we report the effects of EtOH on DA neuron firing activity in a case where Glutamatergic inputs are asynchronous. We report firing rate and burst measure (fraction of spikes within bursts, see methods). On lower panels, we report the voltage trace of the DA neuron for three different EtOH concentrations (0, 1.5 and 3 g/Kg). On the right, we report the same measures but for a case in which glutamatergic units have a certain level of synchronization (f_s_=14%, see Fig.1)

### EtOH boosts DA neuron burstiness mediated by synchronization of Glu inputs

Experimental studies have shown that EtOH increases burstiness in DAergic neuron firing activity (Morikawa et al. 2010). We thus set out to understand how glutamatergic input variability may impact the ETOH effects on the burstiness (measured as percentage of spikes within bursts, SWB, see methods) of the DA neuron and to understand the mechanisms for any eventual differences.

First of all, a visual inspection of simulated DAneuron voltage traces in Fig. 2, shows substantial difference between the two cases of higher and lower Glu input synchronization level. In fact, in the case with low input variability EtOH does not increase DA cell burstiness. In effect, we surmise that under asynchronous glutamatergic drive, EtOH increases only the DA neuron tonic activity. We can clearly see that for higher synchronization levels in Glutamatergic inputs the situation is markedly different. For this case, when EtOH concentrations are relatively high, the DA neuron firing pattern is characterized by the presence of bursts with a high intra-burst firing rate (>12 Hz) and periods of tonic activity with low firing frequency (1-6 Hz). By measuring the number of spikes within burst (SWB) at various EtOH concentrations, we observe a monotonic increase in burstiness for the quasi-synchronous input case as a function of EtOH concentration. We can understand the mechanism as follows: the phasic firing periods of DAergic neuron (bursts) are induced by synchronous AMPA EPSPs, and these are amplified by EtOH through an AMPA conductance increase. At high EtOH concentrations (e.g.3g/Kg), an increase in GABA conductance on DA neurons starts to dominate (e.g. see above for the firing rate effects). Nevertheless, such an increase selectively inhibits DA neuron spikes driven by the tonic AMPA rather then those induced by synchronous AMPA barrages, for which much higher GABA conductance values are necessary. As a result, DAergic neuron spikes that are part of phasic firing periods are maintained also at high EtOH concentration, when GABA conductance is increased, while tonic periods of firing are inhibited. This acts to further boost the fraction of spikes within bursts at EtOH concentrations that decrease the overall average firing rate. In fact, this separable effect on the tonic and phasic DA cell firing by EtOH under partially synchronous glutamatergic input is a prediction of our model.

### Glu input synchronization increases DA neuron sensitivity to EtOH

While we showed that for a partially synchronized glutamatergic input, EtOH robustly increases the burst firing in DA neurons as a function of the dose, we wanted to understand how do different levels of synchronization affect DA neuron sensitivity to EtOH. In order to do so we simulated our model with Glu inputs that had progressively higher fraction of synchronized sources of AMPA EPSC’s while keeping inhibitory GABA and excitatory NMDA inputs fixed.

This unrealistic model (Glutamatergic inputs should activate also NMDA channels) permits us to elucidate the mechanism of cooperation between input synchronization and the increase on AMPA conductance after EtOH. Furthermore, in order to emphasize the net effect of Glu input synchronization, AMPA conductance is changing together with the fraction of synchronous units f_s_, in order to keep constant the average AMPA current received by the DA neuron and investigate the effect of synchronization only.

Our simulations showed that, past a critical synchronization level (as defined by f_s_), both the firing rate and burstiness increase abruptly (Fig. 3 upper panels). In particular, for f_s_ = 10%, DAergic neuron exhibits a transition to bursting dynamics (in the sense of a sharp increase in the burst measure). We further compared the effect of AMPA synchronization across different values of tonic GABA input (Figure 3 lower panels). We find that with increased GABA conductance the qualitative pattern does not change, yet the level of burstiness is significantly increased. In fact, we see generally that above a critical Glu synchronization level, DA cell bursting increases with increasing GABA conductance. Hence our simulations imply amplified sensitivity of DAergic neurons to input synchronization for higher GABA input strength. In other words, the more is the DA neuron inhibited, the more sensitive its burstiness is to synchronous vs. asynchronous excitatory inputs. On the other hand, bottom left panel of Fig. 3 shows that increased GABA conductance monotonically depresses the DA neuron average firing rate, albeit stronger for asynchronous Glu inputs. In Fig 3 lower right panel, we turn to another crucial target of EtOH: the increase of tonic AMPA conductance. We observe that an increase in AMPA conductance on the DA neuron increases its firing activity. Nevertheless, we see that the DA neuron is much more sensitive to such increase in the case of synchronous inputs, while in the asynchronous case the increase of AMPA conductance is much less effective.

**Figure 3:**
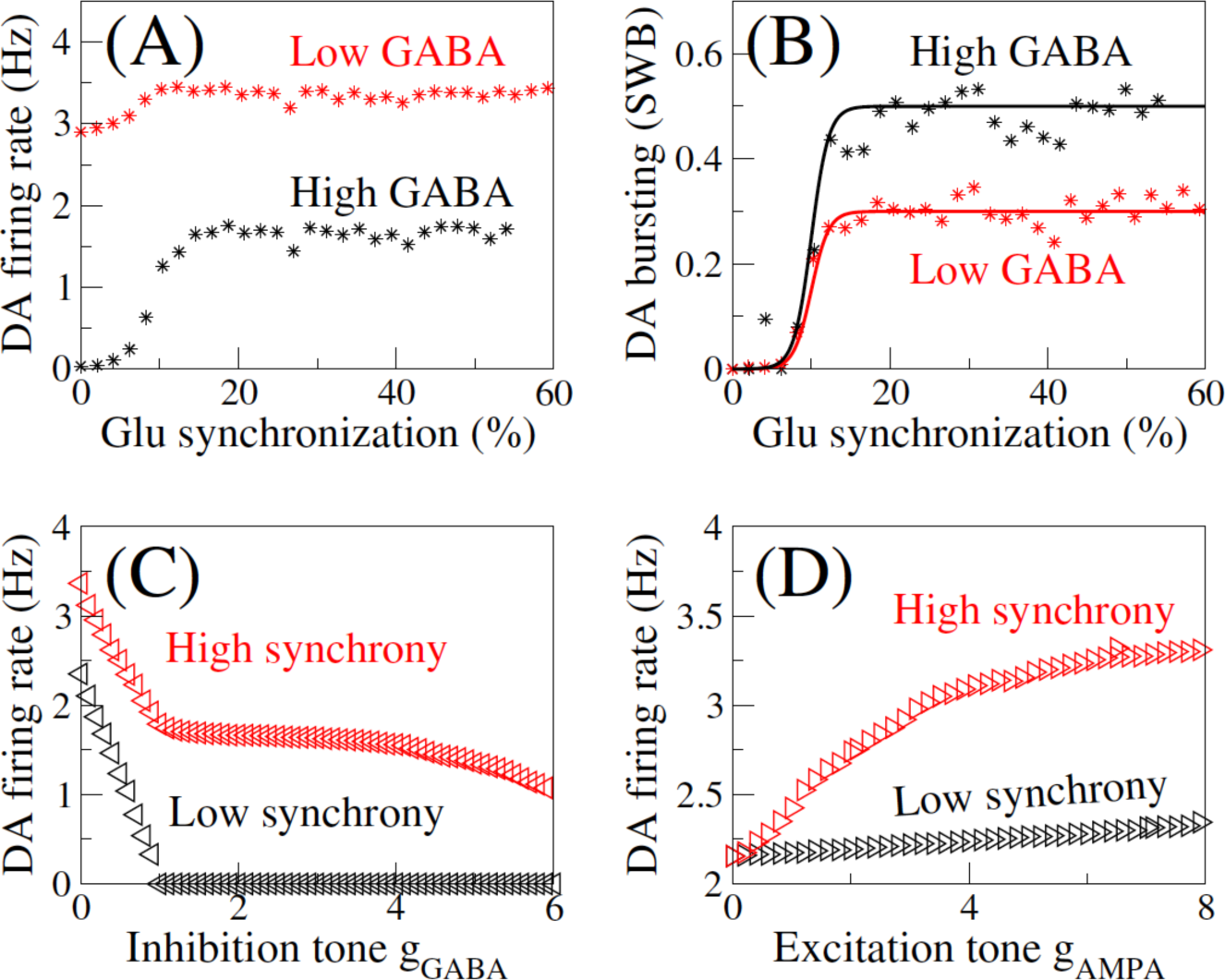
Glu input synchronization, EtOH targets and DA neuron activity. We report here the result of a reduced model: Glu population activity drives only the AMPA input signal to DA neuron, whereas NMDA conductance is set to zero and we consider a tonic GABA IPSC. In top panels: DAergic neuron average firing rate and burst measure respectively as a function of the fraction of synchronous glutamatergic units. For every value of f_s_ the total amount of AMPA EPSC is keeped constant by rescaling appropriately AMPA conductance (equal to 0.03 mS/cm^2^). Lower panels show the effect of increasing tonic GABA (here AMPA conductance is constant g_AMPA_ = 8 mS/cm^2^) and AMPA conductance (here GABA conductance is zero) for two different values of Gluta,matergic unitssynchronization (0 and 14%).

## 2. EtOH effects depends on the average glutameatergic input firing rate

As we saw above, the glutamatergic input synchronization has a significant effect on how sensitive the DA firing patterns are to the EtOH concentration. We reasoned that the average firing rate of the glutamatergic input should also have strong effects on EtOH impact on the DA neuron activity. In order to investigate the correlation of DA neuron activity with glutamatergic rate changes we first investigate the effect of increasing the average firing rate of glutamatergic neurons in an asynchronous case. The main result presented in this section regarding DA neuron average firing arte and is not qualitatively affected by increasing synchronization. We then consider glutamatergic units whose firing pattern follows a Poissonian spike rate with an average changing in time periodically at a certain frequency. It has been shown that cortical regions sending glutamatergic projections to VTA show a prominent coherence with VTA itself and hippocampus at 4 Hz (Fujisawa et al. 2011). By considering such oscillations in inputs we report an interesting interaction between EtOH and oscillatory activity in Glutamatergic inputs: in function of the Glu input rates, EtOH can produce a switch from in phase to antiphase activity of DA neurons with respect to these same inputs (this effect is much more visible for slower Glu input oscillations).

### EtOH inhibits Dopaminergic neuron activity in high frequency Glu input state

Fig.4 shows model simulation results reporting the effects of the Glutamatergic input average firing rate on EtOH influence. We show that, while for relatively low values of ν_Glu_ we observe the U-shaped curve of the DA firing rate as a function of the EtOH dose (as we reported above), in conditions of high ν_Glu_, EtOH has a monotonic inhibitory effect on DA average firing activity. We propose that this is due to an increase in the tonic component of the AMPA conductance during EtOH administration, which, as we have shown before, can silence DA neurons by driving it into depolarization block (Ha and Kuznetsov, 2013). Hence, in our simulations, we see that increasing AMPA conductance speeds up the DA neuron firing but, over a certain threshold value, the DA neuron stops firing. To understand these effects in detail, let us first consider the case of low ν_Glu_. In middle panel of fig 4, we simulate voltage trace of a DA neuron receiving a glutamatergic input at relatively low Glu input rate (10Hz). Upon EtOH application (at 15 seconds), the increase in AMPA conductance has the effect of increasing DA firing activity. On the contrary, in the case case of high ν_Glu_ (lower panel), the DA neuron activity decreases after EtOH injection even if DA neuron is still able to emit some spikes due to noise in inputs. Accordingly, the tonic asynchronous component of Glutamatergic inputs converging into DA neurons can produce a switch in EtOH effects for DA release, i.e. when such component is strong EtOH shows a monotonic inhibitory effect on DA activity.

**Figure 4:**
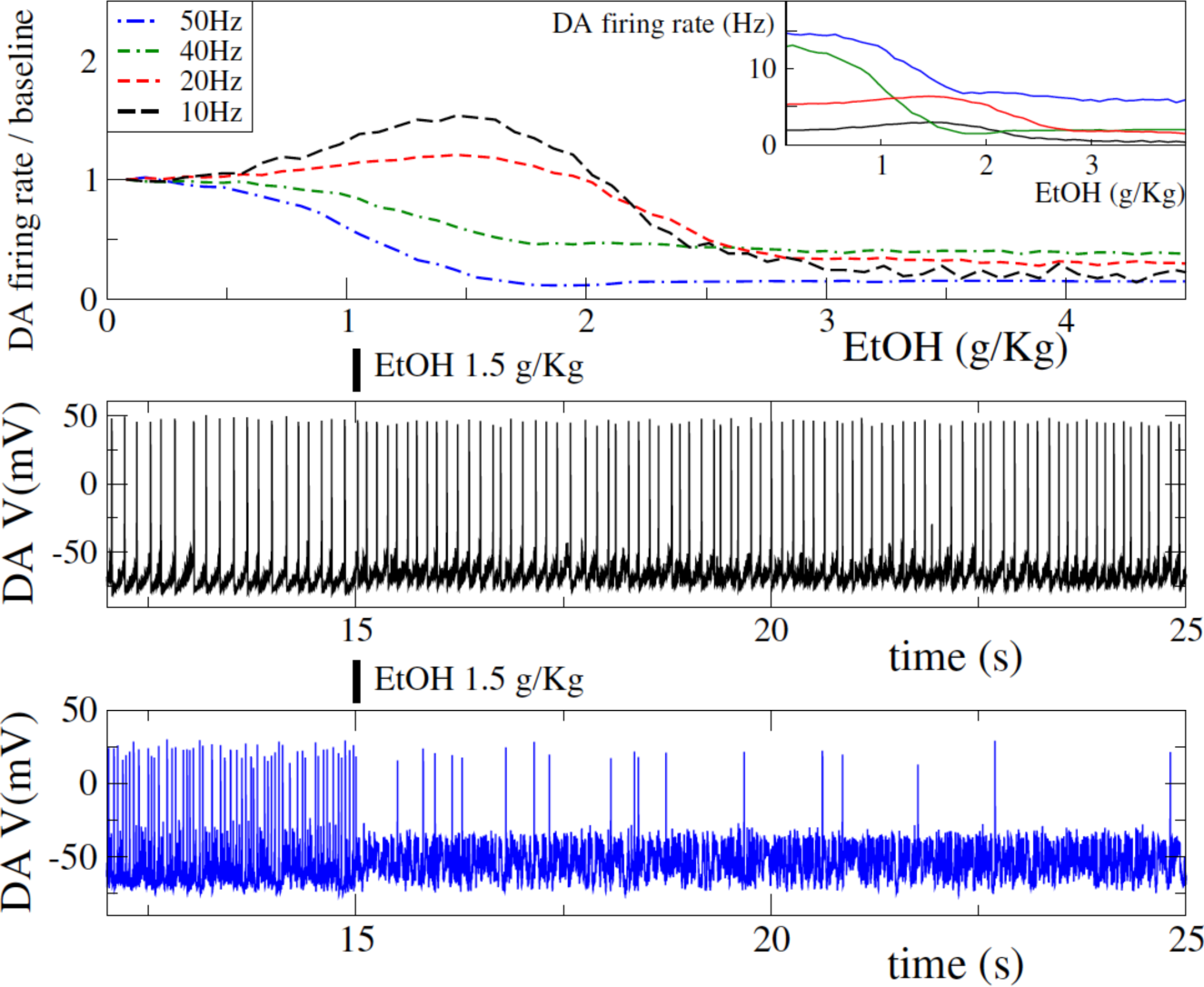
A transition to monotonic inhibition by EtOH at high Glu input activities: EtOH changes input processing of DA neurons in the VTA. We consider here Glu inputs with a firing rate at varouous frequency as reported in the legend. Upper panel shows DA average firing rate with respect to baseline (no EtOH) for the four different Glu frequencies considered. In the inset we show for completeness the real DA neuron firing rate in Hz. Middle panel shows the trace of DA neuron membrane potential for low Glu frequency (10Hz), after 15 seconds EtOH is presented at 1.5g/Kg. Lower panel report the same for higher Glu frequency (50 Hz).

### DAergic neuron and Glutamatergic input coherence

As we see above, EtOH can have a significant effect on the dynamics of the VTA response to incoming Glutamatergic inputs. In previous section the firing rate of Glutamatergic neurons was constant in time. Here, in order to simulate observed oscillations in prototypical cortical inputs to the VTA we consider oscillations in Glutamatergic units firing rate. In particular, we consider theta (4Hz) and delta (1Hz) oscillations. In upper panels of Fig 5 we report the time evolution of Glu units firing rate were we see that, in the case here considered the maximum firing rate is around 50Hz. For such value we know from previous section that DA neuron spikes are blocked from EtOH. In the left panels we report the voltage trace of DA neurons for delta oscillations in Glutamatergic inputs. We observe that DA spikes are suppressed at the peaks of incoming activity when EtOH is present (lower panel). Accordingly while at zero EtOH concentration the DA firing rate is in phase with the Glutamatergic rate (spikes happen around the high level of glutamatergic activity), at high EtOH doses DA activity turns out to be in antiphase with incoming inputs. Hence, the overall conclusion is that higher Glu input levels together with EtOH administration change the relative coherence structure of the DA cell activity: switching it from in phase to out of phase with respect to the cortical activity. This phase/antiphase effect is much clearer considering slower input rate oscillations like delta waves. In fact, in the case of theta oscillations (right column) this effect is less visible and it seems that there is a non trivial relationship between oscillation frequencies of Glutamatergic inputs and the typical time scales of DA neuron intrinsic activity. Nevertheless, even for these fater input frequencies we observe that EtOH changes the coherence between DA and Glutamatergic activity.

**Figure 5:**
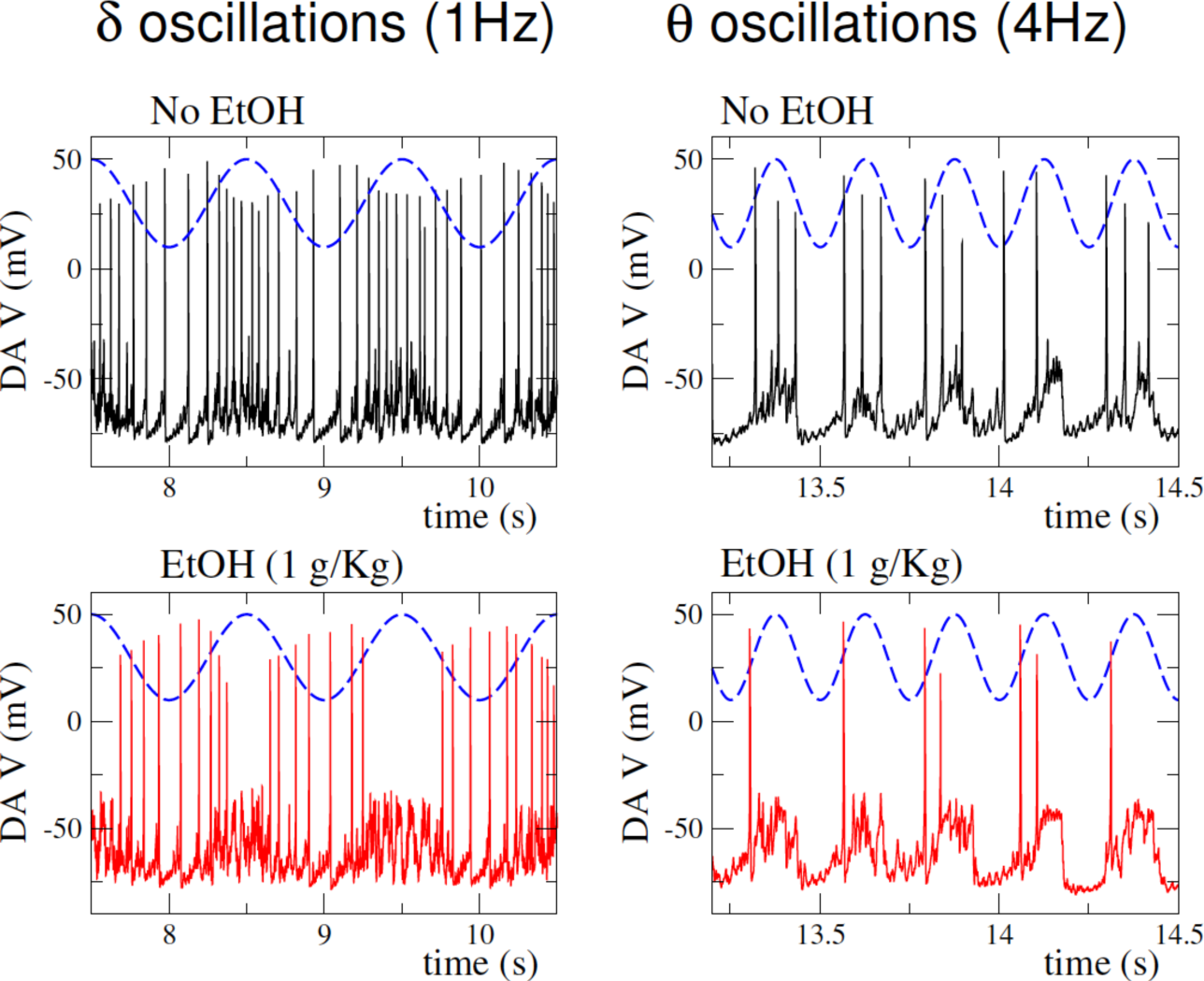
High Glu firing activity guides a transition from in-phase to antiphase DA neuron activity with respect to excitatory inputs. We consider here two different Glutamatergic inputs with a firing rate oscillating like a cosine ν_Glu_=A(cos(2πνt)-1)+ν_0_. Here ν_0_=10Hz, A=20Hz and ν=1Hz or 4Hz for delta (left panels) or theta (right panels) waves. In top panels we report the time course of ν_Glu_. In lower panels we report the corresponding DA membrane voltage trace in case of 0 and 1.5g/Kg EtOH concentration.

## 3. EtOH amplifies high-concentration Dopamine transients through dopamine neuron synchronization

Above we have established how acute EtOH together with glutamatergic inputs leads to complex response patterns from a single DA neuron. Yet we know that the VTA is composed of heterogeneous DA neuron population, whose firing activity regulates the overall DA released. We thus analyze effects of EtOH on DA release from the VTA as a whole by modeling a heterogeneous population of DA neurons in the VTA. We further model spike-based dopamine release for the single DA neurons (see methods section for model details) with the overall DA release modeled as the superposition of these spike-based dopamine transients. Our working hypothesis is that significant increases that are observed in DA concentrations are due to simultaneous firing of DAergic neurons that generate transient peaks of the overall DA release. Hence we hypothesize that synchronization of VTA DA neurons is key to DA release dynamics. Consistent with data (REF) we consider mutual coupling between VTA DA neurons as not being significant, hence it is only common external drive to them that could act as a synchronizing mechanism. If such drive is tonic, i.e. input signal shows little variability (due to a low level of synchronization) DA units with heterogeneous proprieties will not synchronize their firing times. In fact, heterogeneous neuron population receiving time-invariant common inputs would emit spikes depending on their specific proprieties. On the contrary, a transient peak in excitatory input would kick all DA units’ voltage toward the spiking threshold, thus generating a synchronous population burst. In order to test this hypothesis, we simulated 100 DA neurons all receiving common glutamatergic inputs characterized by a certain level of synchronization (f_s_=14%, see Fig.1) and whose heterogeneity was implemented by individual changes in the leakage current picked randomly from a uniform distribution in [0.13:0.23] mS/cm^2^ (see methods). This ensured that these DA neurons differed in their intrinsic excitability. The DA release is calculated as outlined in the Methods. Our simulations indeed show an inverted U-curve for the average DA release (averaged over a sufficiently long time scale to obtain a reliable average measure, i.e. at least one minute in our simulations), with a maximal DA release at an intermadiate EtOH concentration (Fig. 6). The mechanisms standing behind this inverted U shape is the same discussed for the average firing rate of a single DA neuron (see Fig.2). Furthermore, by sampling DA concentration with bins of 180 ms we calculate distributions of the DA levels over an extended period (5 minutes, Fig. XXX). In particular, the figure shows the probablility distributions at low and intermediate EtOH concentrations (around the optimal dose). The distributions are scaled by the mean (which is also EtOH concentration dependent) to emphasize the changes in the width. We see that while in control, DA concentrations are Gaussian distributed around their mean, at the optimal ETOH dose, the distribution shows a large right tail, meaning that there is a significant number of large DA amplitude episodes. Accordingly, the variability in the DA release time trace (Fig. XX) shows a clear increase at these optimal EtOH doses. This underlies another effect of EtOH, i.e. the increased frequency of large deviations from the basal DA level. This effect is moreover elucidated by the time series shown in Fig.6, where we plot DA time traces for different EtOH concentration. We observe that in control condition (EtOH concentration equal to zero) DA stays at an almost constant level with seldom peaks. At higher EtOH concentrations, we observe an increase in the basal level of DA and much more frequent peaks of DA release (middle panel). It is crucial to note that for very high EtOH doses, the DA basal level is reduced, but the peaks are maintained, yielding a lower DA average release with high time fluctuations. What might be the mechanism driving these large amplitude fluctuations in the DA concentration? On the left side of Fig.6, we report the raster for the DA neurons population. We observe that DA peaks of release are driven by synchronous population bursts in DAergic neuron population. In fact, for low EtOH concentration, synchronous inputs received by the VTA are not able to evoke spikes in DAergic units. As a consequence, there are very few DA units firing simultaneously (see the raster plot on left side of Fig.6) and DA release shows little deviation from the baseline. At higher EtOH concentration (1-2g/Kg) the increase in AMPA conductance permits the AMPA EPSCs to be strong enough to evoke synchronous spikes in DAergic neuron population, corresponding to high DA concentration transients (see middle panels, 2g/Kg). The baseline DA release is also increased in EtOH. The changes in DA neurons intrinsic activity caused by EtOH are responsible for this as they increase firing activity throughout the simulation including asynchronous intervals.

**Figure 6.**
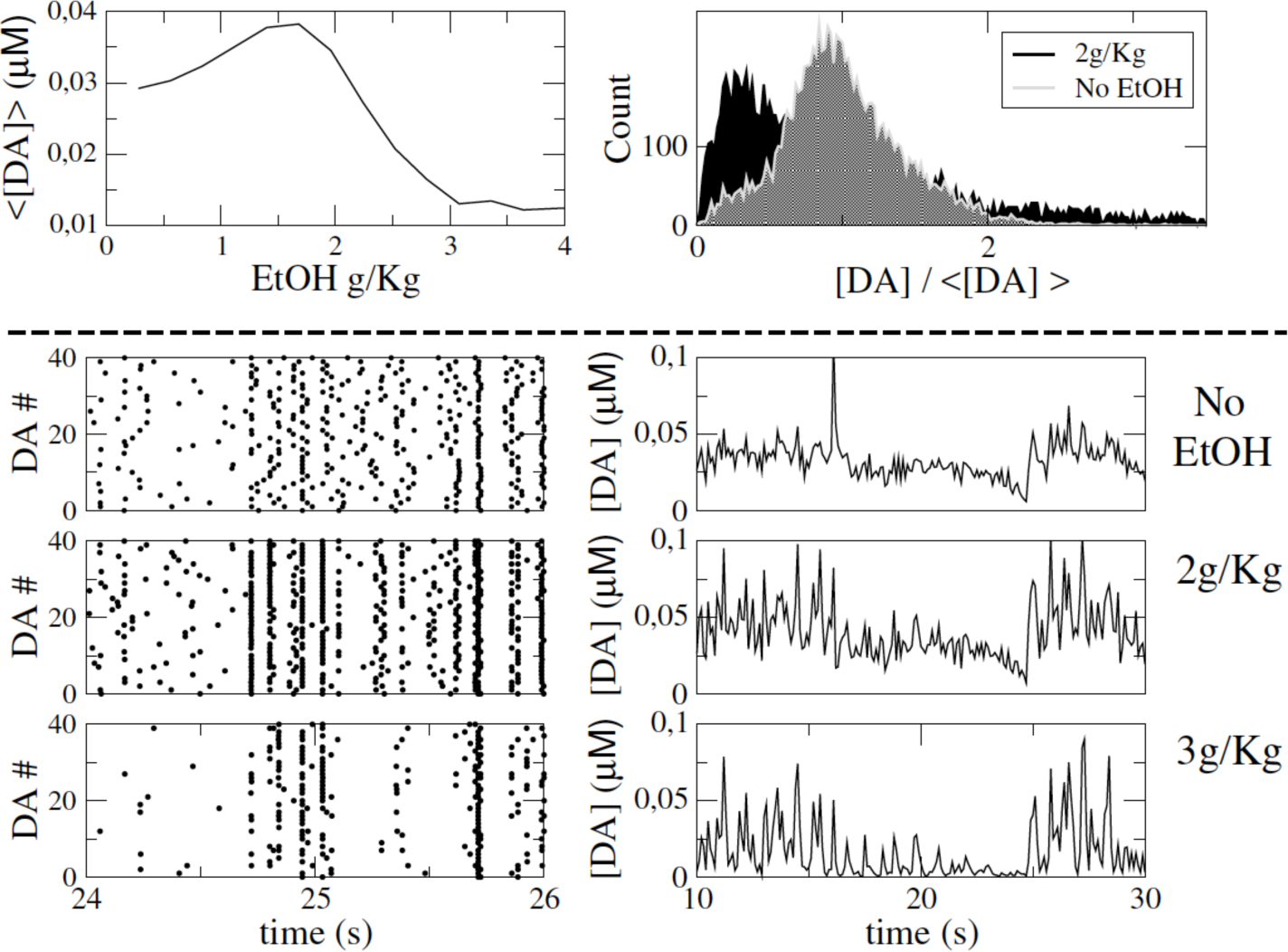
EtOH yields transient release of Dopamine from VTA by enhancing synchronization between DAergic units. Top-left: average (over one minute time scale) DA release by the population of 100 DA neurons at various EtOH concnetration. (B) Histogram of DA concentration (rescaled by its average) for two EtOH concentration as reported in the legend. Left panels show the raster plot of a subpopulation of DA neurons and the corresponding time trace of DA release at various EtOH concentrations (0,2,3 g/Kg). Glutamatergic inputs are quasi-synchronous (f_s_=14%) and ν_Glu_ =14 Hz.

Eventually, at EtOH doses above the optimum, EtOH-induced increase in GABA input inhibits tonic background asynchronous activity of the DA population, thus reducing basal DA levels. Nevertheless, the synchronous population bursts of DA neurons are only little affected by EtOH (as described in previous section, see Fig. 2 and 3). Accordingly, at high EtOH doses DA release is characterized by a lower baseline level but still amplified transient DA peaks.

Again, variability and synchronization in input structure turns out to be a crucial ingredient for evoking transient peaks in DA release. In cooperation with EtOH influence, such correlations in input structure have a double effect. On one side, they permit EtOH to evoke phasic firing episodes in single DA units (see Fig.2) and, on the other side, to synchronize these phasic episodes, thus yielding transient peaks in DA release from VTA.

## Discussion

Taking advantage of modeling techniques, we built up a circuit model that simulates the effects of various EtOH concentration on the dynamics of VTA Dopamine neurons and their Dopamine release. In particular, we focused on the way external Glutamatergic inputs are processed by VTA and then converted into Dopamine release, as a meaningful output projecting to NAc regulating reward proprieties and, ultimately, addiction.

By following experimental measures we derived EtOH dependent values of conductance of HCN channels and GIRK current able to reproduce input independent effects of EtOH observed in vitro (see method section). We then calibrated EtOH effects on DA neuron input conductances, i.e. AMPA and GABA.

Such a model is able to reproduce EtOH effects at the level of single DAergic neuron and at the level of their collective activity, i.e. their DA release from VTA to the Nucleus Accumbens (NAc).

In particular, the model correctly predicts the concentration dependent effect of EtOH on DA neuron firing pattern, i.e. an increase in phasic (bursting) activity and an increase of average firing rate followed by a decrease at high EtOH concentrations.

The mechanism proposed is based on the interplay of EtOH targets and Glutmatergic input synchronization. In particular, synchronized inputs generate DA neuron phasic activity periods, whose magnitude and frequency of appearance is increased from AMPA conductance increase from EtOH. Furthermore, the decrease of DA activity is due to delayed activation of GABA conductance at higher EtOH concentration, inhibiting tonic DA neuron period of firing.

At the level of population activity, the model reproduces an increase in transient peacks of Dopamine release, that has been reported to be a robust and distinctive effect of EtOH and other drugs.

The mechanism here proposed is based synchronous inputs drive the synchronization of these phasic periods between different DA neurons, that would be asynchronous otherwise due to their heterogeneity. Synchronization of DAergic neurons after reward events has been reported in the midbrain by Joshua et al. (2009). Our model predicts this to be the case also in the VTA after acute EtOH injection and, more generally, as a result of addictive drugs.

The mechanism of synchronization relies on EtOH-mediated enhancement of AMPAR current that increases the probability to evoke spikes in response to synchronous Glu pulses. Complementary mechanism of EtOH-mediated synchronization of DA neurons in case they receive noisy correlated Glu inputs is discussed in our previous work (Morozova et al 2016). We showed that EtOH-mediated enhancement of Ih and/or AMPA currents produce higher levels of synchronization between heterogeneous DA neurons by switching their excitability type to type II. The responses of type II neurons to noisy inputs are more reliable that of type I neurons (Gutkin et al., 2005; Galan et al., 2007). We showed that synchronous activity of DA neurons produces higher DA transients. This result further confirms the role of synchrony in EtOH-evoked enhancement of DA transients.

Moreover, the model predicts a different effect of EtOH on VTA DA neurons activity for different Glutamatergic input rates.

We report in fact that for high Glutamatergic input average rate EtOH has a monotonic inhibitory effect on DA neurons, while for lower input rate it shows the expected increase followed by a decrease at high EtOH concentration. This result, as the necessity of a certain level of synchronization, predicts an input dependent effect of EtOH on DA release. In particular, the way EtOH affects DA release is dependent on the actual state of Gutamatergic inputs, its synchronization and average rate.

It is known that VTA receives Glutamatergic afferents from many regions, like the PFC.

Rephrasing our result the model predicts different DA response to EtOH injection for different PFC activity state. This prediction could be verified by measuring DA concentration in NAc under different concentration levels or under direct stimulation of PFC.

Furthermore, by considering oscillatory firing rate in Glutamatergic inputs (in accordance with oscillations reported by Fujishawa et al. 2011), our model predicts that Ethanol produces a switch between in phase to in antiphase DAergic activity with respect to Glutamatergic input. On the same line with the previous one this prediction could be tested through direct stimulation of PFC after Ethanol injection.

Actually, Ethanol itself affects the firing rate of PFC, specifically by decreasing it (Tu et al. 2007). In the view of our results this would have the effect of decreasing DA neuron firing rate. Nevertheless, a direct outcome of this model is that the average measures are not the only properties of afferent signals that affect DA release. In particular, synchronization and fluctuations if Glutamatergic neurons rate are crucial ingredients to increase DA neuron phasic activity and, ultimately, DA release transients. Accordingly, an interesting test to be performed experimentally is if, during in vivo conditions where DA neuron bursting has been showed to be increased by EtOH like other addictive drugs (Covey et al. 2014), PFC activity increases its coherence and deviations from average.

Another possible mechanism is based on GABA neurons activity in the VTA. It has been shown that GABA neurons average rate decreases after EtOH injection (Stobbs et al. 2014). This has the effect of increasing DA neurons activity through dishinibition. Accordingly, the decrease of afferent excitation to the VTA could increase DA neuron firing rate through this dishinibition mechanism. In our model this is not the case as the effect of the direct decrease of excitation to DAergic neuron is stronger then dishinibition.

Nevertheless, even hypothesising this to be the case (through some model parameter arrangement), the decrease of average PFC rate would yield an increase in tonic activity of DA neurons and not in their phasic periods, responsible for DA transients in NAc. Accordingly, based on these observations and our modelling results, we predict that EtOH (and it might be the case for other drugs) should affect coherence and synchronization proprieties in cortical regions. Furthermore, an increase in synchronization in inputs received by VTA would synchronize also GABA neurons, which have been reported to boost phasic DA neuron activity when they synchronize with each others (Morozova et al. J. Neurophysiology 2016).

In conclusion, the circuit model here presented is able, from the one side, to reproduce the observed effect of Ethanol on DA neurons activity and their DA release and, on the other side, to make testable prediction. In particular, synchronization of DA neurons after EtOH injection is proposed as the main mechanism for generation of transient peaks of DA concentration in NAc. Moreover, synchronization in cortical VTA afferents is proposed to be a fundamental ingredient for increase in DA transients after EtOH injection and, more generally, cortical neurons activity state can drive opposite effects of EtOH on DA response.

## Methods

We describe here the circuit model adopted for modeling the VTA, its input and its output.

### Inputs: Glutamatergic population

We use a putative model of the pyramidal cell population in the PFC, generating spike trains of glutamatergic neurons. Such firing pattern is characterized by two main parameters: the level of activity and the correlation between units. Every unit emits spikes according to a Poissonian process with an average firing rate ν_Glu_. Regarding the units correlation we impose that, during certain time windows, a fraction f_s_ of units emits spikes almost synchronously. In particular, we force them to be randomly confined in a time interval of 5ms. The time duration of synchronous and asynchronous time windows is extracted from a poissonian distribution with the same T_s_=4s. In the raster plot of Fig.1A we show an example with ν_Glu_=4Hz, f_s_=0 and 14% (unit number from 0 to 7, we consider N_E_=50 Glutamatergic units).

In all our simulations the number of units is fixed at N_E_=50.

Changing f_s_ yields a change in fluctuations of Glutamatergic units count number CN(t), i.e. normalized (with respect to the number of units) number of spikes of the all population in time intervals of 50ms (as reported in Fig. 1A).

The excitatory activity in this population will furnish the input to the VTA through AMPA and NMDA receptors located on both GABA and DA neurons. Hence we will convert the spike trains into synaptic inputs keeping track of PFC to VTA input convergence.

The dynamics of AMPA and NMDA receptor is the same in both GABA-ergic and DA-ergic neurons. What changes in the two neuron types is their maximal AMPA and NMDA conductance. The AMPA current flowing into the DA-ergic neuron (calling V(t) its membrane voltage) is I_AMPA,DA_=g_DA,AMPA_p_AMPA_(t)(E_AMPA_-V(t)), where g_DA,AMPA_ is the maximal conductance of AMPA receptor on DA-ergic neurons and p_AMPA_(t) is the gating variable that depends on Glutamatergic units spike train. We use the same notation for NMDA, I_NMDA,DA_=g_DA,NMDA_p_NMDA_(t)(E_NMDA_-V(t)). Let us define q_i_ the activation variable of unit *i* in the population of Glutamatergic units. Unit *i* is active, i.e. with activation variable q_i_=1 (inactive q_i_=0), for an amount of time equal to 1 ms around the spiking time. The number of inputs that the neuron receives at time t, say Q_i_(t), is the sum of q_i_(t) over all the N_E_ Glu units. Not every input is able to open completely AMPA or NMDA channel, and we use a sigmoidal function to obtain the variable j= 1/(1+e^-(Q-9)/1.3)^ that is the fraction of open AMPA and NMDA channels. Then, in order to obtain the gating variable of AMPA and NMDA channels we use the following equations:

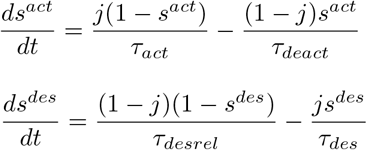

The gating variable p_AMPA_(t) is equal to the product of desensitisation and activation variables, i.e. p_AMPA_(t)=s^act^s^des^, while p_NMDA_(t) is equal to the activation variable. The dynamical evolution of NMDA and AMPA channels differs also for time scale of activation and deactivation, where the last one is much longer in NMDA channels. The input j is a function of the spiking pattern of Glutamatergic pattern, and thus depends on the average firing rate and their synchronization level f_s_.

### VTA circuit: GABAergic population

Voltage dynamics of each GABA neuron is described by Wang-Buszaki model (see appendix I) of fast spiking neurons with a parameter distribution to reflect heterogeneity and coupling will be described by the typical electrical coupling function (Wang and Buzsaki, (1996) and Kepler et al., (1990)).

Each neuron i is coupled with all the other neurons j in the population with the typical electrical coupling I_el_=g_el_(v_i_-v_j_). This is a heterogeneous population (Margolis et al., 2012) that fires regularly (e.g. pacemaking) at high frequencies. Experimental data suggests that the range of firing rates of recorded VTA GABA neurons is very broad with the mean of 19Hz (Steffensen et al., 1998). The differences in frequencies are modeled by changing the leak conductance g_lg_, according to g_lg_=0.05+0.05*(*rnd*-0.5), where g_lg_=0.05 corresponds to frequency of 17 Hz. GABA neurons are capable of firing with much higher frequencies in response to excitatory synaptic inputs, modeled by NMDA and AMPA currents. The activity of each GABA neuron contributes to a GABA current flowing into the DA neuron. Every time a GABA neurons fires, a certain amount of neurotransmitter is released, which contributes to activation of the gating variable according to the following equation (where we indicate *v*_i_ the voltage of neuron i_th_):

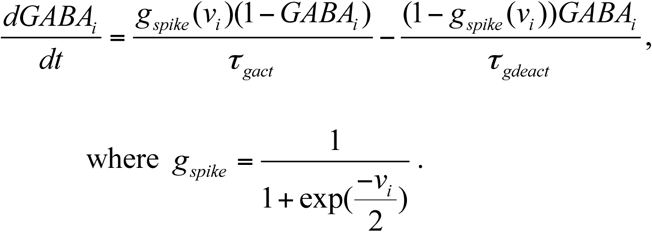

We consider a population of 50 GABA neurons. Every DA neuron receives different GABA inputs according to the dynamics of a subpopulation of M=10 randomly chosen GABA units neuron.

In particular, the gating GABA variable of a specific DA neuron is the average over M randomly chose units in the GABA population:

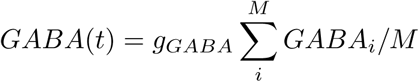

### VTA circuit: DA-ergic population

The model of the DA neuron is a conductance-based one-compartmental model:

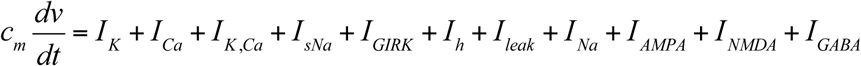

Where *v* is the voltage and the first seven currents represent intrinsic currents: a calcium current I_Ca_, a calcium-dependent potassium current I_K,Ca_, a potassium current I_K_, a subthreshold sodium current I_sNa_, a sodium current I_Na_ and a leak current I_leak_. AMPA and NMDA receptor currents (I_AMPA_ and I_NMDA_, respectively) model excitatory inputs and GABA receptor current (I_GABA_) models inhibitory inputs. The first subgroup of currents mostly contributes to pacemaking mechanisms of DA neuron, while synaptic inputs produce bursts and pauses. In Appendix 1 we write the details of such currents, their dynamics and voltage dependence while in table 1 we report the parameters, chosen according to Morozova et al (2016). We model a heterogeneous population of ten DA neurons, each one receiving inputs from different GABA neurons and with different leakage currents extracted randomly in the interval [0.13:0.23] mS/cm^2^.

### Output: DA release

Every DA neuron releases a certain amount of Dopamine depending on its spike train {t_s_}. The following equation relates the spiking times of DA-ergic neuron to the amount of dopamine released:

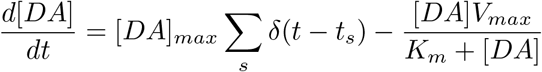

where δ(t-t_s_) is a delta function that increases of a finite fraction [DA]_max_ the dopamine concentration [DA].

The second term represents DA uptake described by Michaelis-Menten equation, where V_max_=0.004 µM/ms is the maximal rate of uptake by a transporter and K_m_=0.2 µM is the affinity of the transporter for dopamine. The total amount of Dopamine released by the system is the linear sum of the dopamine released by each DA neuron.

### Ethanol effects: EtOH concentration and model parameters change

Ethanol targets we take into account can be separated in two kinds: intrinsic and input targets.

*DA intrinsic currents:* Ethanol effects on intrinsic activity are the increase in GIRK current conductance and in HCN channels conductance. We calibrate Ethanol effects in a way that high EtOH concentration reproduces the results shown in McDaid et al. (2007). Ethanol increases of around 150% DA-ergic neuron firing rate in vitro and, in absence of I_h_ current, Ethanol produces a derease in DA-ergic neuron firing rate. In panel a) of Fig.7 we plot the effects of increase in g_h_ and g_GIRK_. The increase in I_h_ conductance has a double effect, i.e. at very high values of g_h_ DA-ergic neuron stops firing while for low values, an increase of in I_h_ conductance increases DA-ergic neuron activity. In order to reproduce the result of Mc Daid et al. we impose EtOH to move such parameters as shown by the arrow in the Fig.7a. By doing so, we can see in panel b) how the effects we were looking for are well captured. Through this calibration we obtain control and high EtOH concentration values of intrinsic currents in DA-ergic neuron. Then, we suppose such parameter to depend on EtOH concentration through a sigmoidal function, and in panel c) of Fig.7 we show how g_h_ and g_GIRK_ depend on EtOh concentration [EtOH].

**Figure 7.**
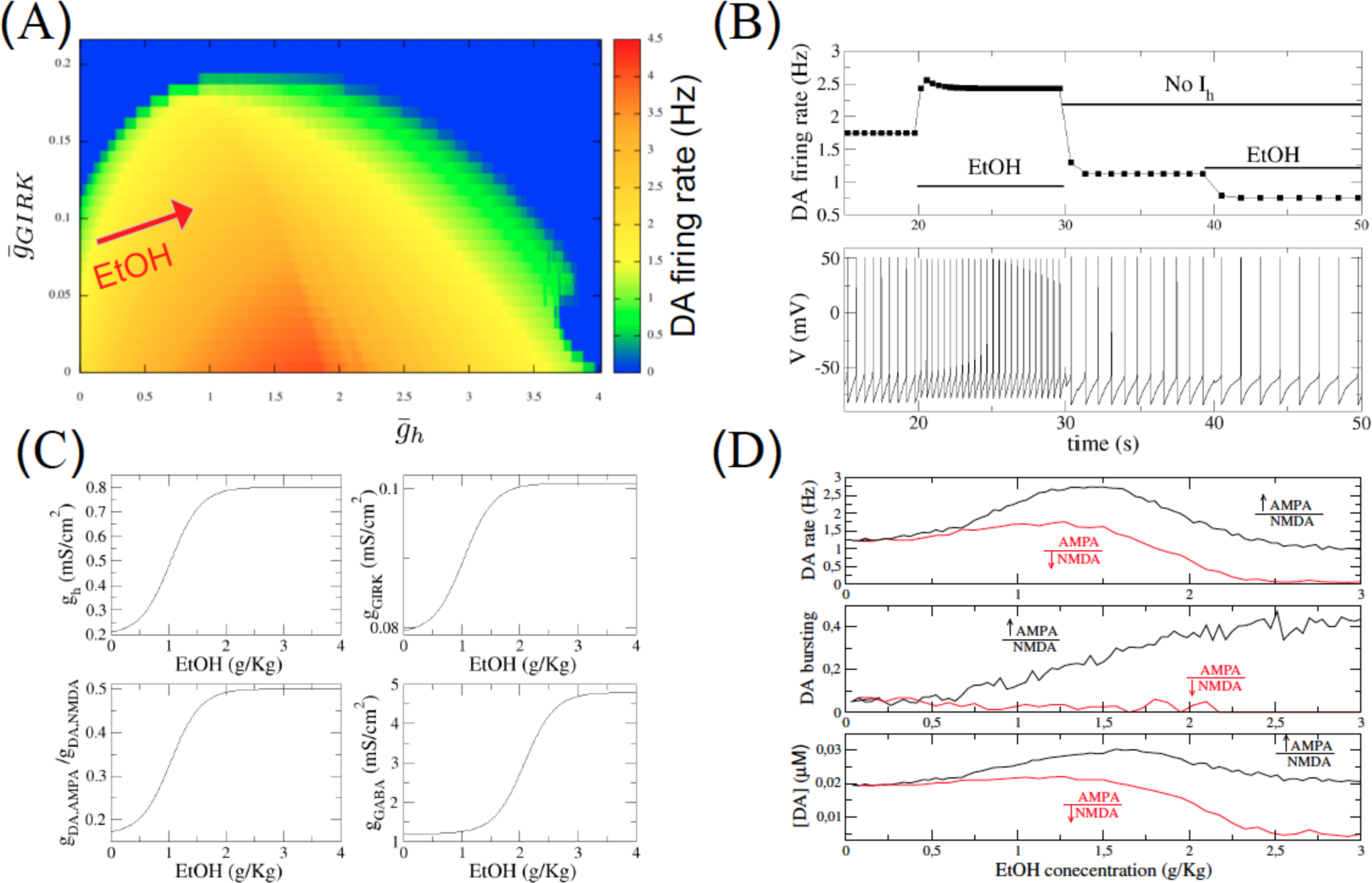
EtOH model calibration. (A) Combined effect of increasing g_h_ and g_GIRK_ on DA neuron firing rate. In accordance with experimental observations at high values of g_h_ DA firing rate decreases. The red arrow represents our parameter setup for EtOH effects on DA neuron intrinsic activity. (B) The model reproduces the observation that EtOH has inhibitory effect when g_h_=0. (C) Parameters dependence on EtOH concentration. (D)) DA neuron average firing rate, bursting and DA release from the population of DA neurons in function of EtOH concentration. Black curves refear to the case in which we increase AMPA conductance to increase AMPA/NMDA ratio as in Saal et al. while red ones refear to the case in which we decrease NMDA concuctance to obtain the same ratio increase (Input parameters are the same of Fig.1).

Regarding EtOH effects on VTA input we take into account an increase in AMPA/NMDA ratio and an increase in GABA conductance of DA-ergic neurons. To calibrate AMPA/NMDA ratio increase we follow the observations of Saal et al. (2003) where the ratio between AMPA and NMDA EPSP was increased by a factor around 1.5 with a concentration [EtOH] =0.02g/Kg. As we are considering concentration up to 2g/Kg in order to reproduce EtOH dose concentration effects as reported by Mereu et al. (1984) we suppose an increase up to three times for very high EtOH doses. Accordingly, in panel c) of Fig.7 we plot the variation of gDA,AMPA/gDA,NMDA in function of Ethanol concentration [EtOH]. Again, in the view of reproducing an increase in DA neuron average firing rate and bursting and in accordance with experimental observations by Ding et al. (2012) we model the increase in the ratio by the increase of AMPA conductance. For the sake of completeness we show in Fig. 7d that the opposite choice (decreasing NMDA conductance) would yield a purely inhibitory of EtOH on DA neuron activity and DA release.

Regarding EtOH effects on g_GABA_ we model an increase in GABA conductance of DA neurons as it has been showed by Theile et al. (2008) that Ethanol enhances GABA transmission in DA neurons in the VTA. We calibrate it in a way that our model is able to reproduce the result by Mereu et al. (1984), where it is shown that high EtOH concentrations ([EtOH]>2g/Kg) produces a decrease in DA neurons firing rate with respect to baseline level. We choose the function g_GABA_([EtOH]) (see panel c) of Fig.7) in a way to reproduce this effect and to observe a reasonable (1-4Hz) firing rate of DA neuron in control condition ([EtOH]=0g/Kg), as shown in panel d) of Fig.7.

### Spike within bursts SWB

To analyze the bursting features of DAergic neuron we calculate the percentage of spikes within a burst (SWB). This measure is calculated defined on the basis of 5 minutes of simulation time with a minimum 200 spikes in this time interval. Bursts are identified as discrete events consisting of a sequence of spikes with burst onset defined by two consecutive spikes within an interval less than 80 msec, and burst termination defined by an interspike interval greater than 160 msec (Grace and Bunney, 1984b). The SWB was calculated as a number of spikes within bursts divided by the total number of spikes.

**Table:** Model parameters

| Parameter | Description | Value |
| --- | --- | --- |
| $C_m$ | Membrane capacitance of DA and GABA neurons | $1\mu F / cm^2$ |
| $\bar{g}_K$ | Maximal potassium conductance on DA neuron | $1mS / cm^2$ |
| $\bar{g}_{Ca}$ | Maximal calcium conductance on DA neuron | $2.5mS / cm^2$ |
| $\bar{g}_{KCa}$ | Maximal calcium-dependent potassium conductance on DA neuron | $7.8mS / cm^2$ |
| $\bar{g}_{sNa}$ | Maximal subthreshold sodium conductance on DA neuron | $0.13mS / cm^2$ |
| $g_l$ | Leak conductance on DA neuron | $0.18mS / cm^2$ |
| $\bar{g}_{Na}$ | Maximal sodium conductance on DA neuron | $50mS / cm^2$ |
| $\bar{g}_{Nag}$ | Maximal sodium conductance on GABA neuron | $22mS / cm^2$ |
| $\bar{g}_{Kg}$ | Maximal potassium conductance on GABA neuron | $7mS / cm^2$ |
| $\bar{g}_{AMPA,GABA}$ | AMPA conductance on GABA neurons | $0.8mS / cm^2$ |
| $\bar{g}_{NMDA,GABA}$ | NMDA conductance on GABA neurons | $0.5mS / cm^2$ |
| $\bar{g}_{Gap}$ | Gap junction coupling between GABA neurons | $0.02mS / cm^2$ |
| $\bar{g}_{NMDA}$ | Maximal NMDA conductance on DA neuron | $18mS / cm^2$ |
| $g_h$ | Maximan HCN conductance on DA neuron | varied |
| $g_{AMPA}$ | AMPA conductance on DA neurons | varied |
| $g_{GABA}$ | GABA conductance on DA neurons | varied |
| $\bar{g}_{GIRK}$ | Maximal GIRK conductance on DA neuron | varied |
| $E_h$ | HCN reversal potential on DA neuron | $-20mV$ |
| $E_K$ | Potassium reversal potential on DA and GABA neurons | $-90mV$ |
| $E_{Ca}$ | Calcium reversal potential on DA neuron | $50mV$ |
| $E_{Na}$ | Sodium reversal potential on DA and GABA neurons | $55mV$ |
| $E_l$ | Leak reversal potential on DA neuron | $-35mV$ |
| $E_{lg}$ | Leak reversal potential on GABA neuron | $-51mV$ |
| $E_{NMDA}$ | NMDA reversal potential on DA and GABA neurons | $0mV$ |
| $E_{AMPA}$ | AMPA reversal potential on DA and GABA neurons | $0mV$ |
| $E_{GABA}$ | GABA reversal potential on DA neuron | $-90mV$ |
| $\tau_{aact}$ | AMPA receptor activation time on DA and GABA neurons | $1ms$ |
| $\tau_{adeact}$ | AMPA receptor deactivation time on DA and GABA neurons | $1.6ms$ |
| $\tau_{nact}$ | NMDA receptor activation time on DA and GABA neurons | $7ms$ |
| $\tau_{ndeact}$ | NMDA receptor deactivation time on DA and GABA neurons | $170ms$ |
| $\tau_{gact}$ | GABA receptor activation time on DA and GABA neurons | 0.08ms |
| $\tau_{gdeact}$ | GABA receptor deactivation time on DA and GABA neurons | 10ms |
| $I_{app}$ | Applied current on DA and GABA neurons | 0 |

## Appendix 1

### A1.1 DA neuron model details

Two Calcium currents are present in DA neuron model: an L-type voltage-dependent calcium current *I_Ca_* = *g_Ca_*(*E_Ca_* − *v*) and an SK-type calcium-dependent potassium current *I_K,Ca_* = *g_K,Ca_*(*E_K,Ca_* − *v*). Gating of the calcium current is instantaneous (Wilson and Callaway, 2000; Helton et al., 2005) and described by the function

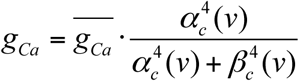

Calibration of the calcium gating function reflects an activation threshold of an L-type current, which is significantly lower in DA neurons than in other neurons, ∼ -50mV (Wilson and Callaway, 2000; Durante et al., 2004). Calcium enters the cell predominantly via the L-type calcium channel. Contribution due to the NMDA channel is minor (Oster and Gutkin, 2011). A large influx of Ca^2+^ leads to activation of the SK current, which contributes to repolarization as well as after hypolarization of the DA cell. Dependence of the SK current (*I_K,Ca_*) on calcium concentration is modeled as follows (Kohler et al., 1996)

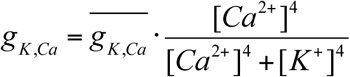

Calcium concentration varies according to the following equation:

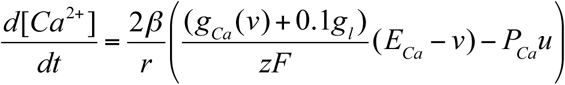

This equation represents balance between Ca^2+^ entry via the L channel and a Ca^2+^ component of the leak current, and Ca^2+^ removal via a pump. In the calcium equation, *β* is the calcium buffering coefficient, i.e. the ratio of free to total calcium, *r* is the radius of the compartment, *z* is the valence of calcium, and *F* is Faraday’s constant. *P_Ca_* represents the maximum rate of calcium removal through the pump.

The neuron is repolarized by the activation of a large family of voltage-gated potassium channels. The model contains voltage-dependent K^+^ current *I_K_* = *g_K_*(*E_K_* − *v*). Conductance of this current is given by a Boltzmann function:

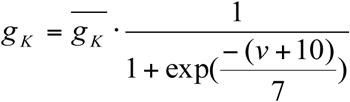

The DA neuron expresses voltage gated sodium channels that carry a large transient current during action potentials (the spike-producing sodium current and a noninactivating current present at subthreshold voltages (a subthreshold sodium current *I_sNa_* = *g_sNa_*(*E_Na_* − *v*)). Even though the persistent subthreshold sodium current is much smaller than the transient spike-producing current, it influences the firing pattern and the frequency of the DA neuron by contributing to depolarization below the spike threshold. We modeled the voltage dependence of the subthreshold sodium current as follows

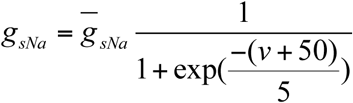

The kinetics and the voltage dependence of the subthreshold sodium current were taken from Carter et al. (2012). The spike-producing sodium current has the form of Hodgkin-Huxley model *I_Na_* = *g_Na_m*^3^*h*(*E_Na_* − *v*), where sodium activation and inactivation channels obey the following differential equations:

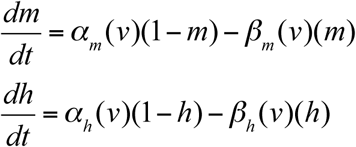

where the rate cosntatns α_x_ and β_x_ have the following form:

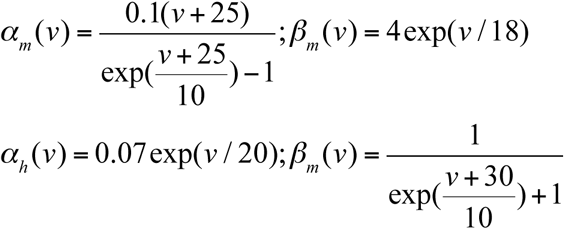

The leak current (*I_l_* = *g_l_*(*E_l_* − *v*)) in the model has the reversal potential of -35 mV, which is higher than in the majority of neuron types. In DA neurons, several types of depolarizing, nonselective cation currents are expressed, which likely contribute to depolarization during interspike intervals. The GIRK current *I_GIRK_* = *g_GIRK_*(*E_K_* − *v*) is modeled as a leak potassium current. The HCN channel *I_h_* = *g_h_q*(*E_h_* − *v*) depends from the activation variable q that obeys to the following dfferential equation:

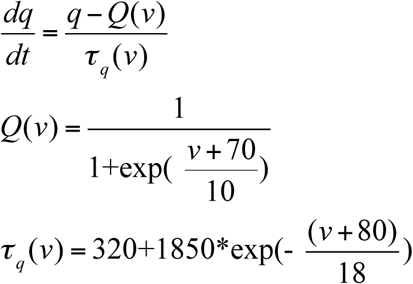

where, in the last equation, quantities are measured in ms.

### A1.2 GABA neurons model

GABA neuron dynamics obeys the following differential equations:

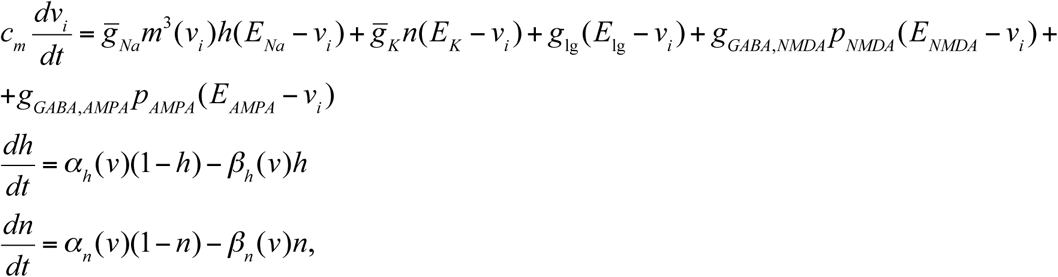

where *v_i_* is the voltage of cell i and the activation variable equations are completed by the following relations:

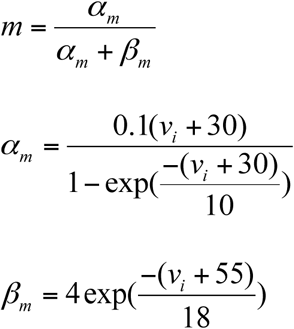

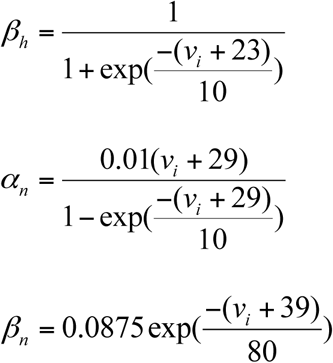

